# Building Inclusive Leadership: A Biosciences Approach

**DOI:** 10.1101/2024.12.09.626597

**Authors:** Emily May Armstrong, Lydia Bach, Philip Robinson, Helen Walden, Nathan Woodling, Matt Jones

## Abstract

One-size-fits-all Equity, Diversity, and Inclusion (EDI) strategies and provisioning are no longer suitable for a growing and diversifying higher education landscape; as each research discipline faces its own unique challenges to inclusion. High impact research needs reliable, decisive, and holistic approaches to leadership, but managerial and leadership skills are often considered an ‘add-on’ as opposed to a necessity in modern research environments. An estimated 70% of EDI initiatives in UK Higher Education Institutions fail to flourish (Marchiondo *et al*. 2023) – a combinatory effect of conservative approaches to change, cultural adaptation, and departmental stability.

We developed an Inclusive Leadership Programme to empower bioscientists to work autonomously and inclusively. Using a co-creation approach, the Inclusive Leadership Programme collectively defined inclusive leadership traits to create a resource. The interactive resource shares inclusive approaches to communication, feedback, recognition, and development – prioritising the experiences of those typically excluded from leadership conversations. By embedding a self-reflective scoring system, we share changes in approaches to inclusivity. After using the resource, self-perception normalises in distribution, indicating the resource provides a baseline for best practice, and space to understand personal approaches to inclusive practice in biosciences.

Ultimately, the Inclusive Leadership Programme and Resource offers a template for other biological research, learning, and teaching environments to build their own tailored approach to integrating inclusivity and empowering all members to work as autonomous and inclusive leaders, regardless of seniority.

## Introduction: The need for proactive inclusion

A robust, effective, and relevant Equity, Diversity, and Inclusion (EDI) culture is key to organisational success in the global Higher Education (HE) sector. Embedding effective EDI approaches within research is now an awarding requirement for the UK Research and Innovation grants from 2023 onward (UKRI, 2023). An engaged approach to inclusion is becoming a requirement to produce meaningful and relevant research within the biosciences (Davies *et al*. 2021). This is exemplified in sector-wide efforts to improve gender disparity in leadership, decolonise the Euro-centric curriculum (Winter *et al*. 2024), challenge the awarding gap (Davies *et al*. 2021), and tackle under-representation of global ethnic majority researchers (Hughes *et al*. 2023), disabled researchers, and LGBTQIA+ researchers in biosciences (Helman *et al*. 2020).

Despite progress being made to improve disparities, numerable challenges remain. Upwards of 70% of EDI-oriented research-culture projects fail to create meaningful changes, attributed to in part by institutionally conservative approaches to balancing cultural adaptation and stability (Hodgins *et al*. 2022). Compounding these factors further, Marchiondo *et al*. (2023) highlights the impacts of EDI programmes can be further limited by a lack of engagement from leadership and senior institutional members, regardless of seniority, job role, or title. This lack of engagement is sometimes attributed to academic promotion and reward criteria that prioritise perceived academic excellence, thus deprioritising formal managerial training and experience. This presents a difficult trade-off, where organisations must facilitate and promote autonomy, while balancing unified approaches to leadership.

Department organisation, combined with EDI resource availability and relevance, present additional challenges to integrating inclusivity. As HE diversity increases, HE institutions are creating and engaging with bespoke and collaboratively-driven EDI resources, reflecting the needs of a given cohort (AdvanceHE, 2024). Additionally, as each research discipline faces different inequalities (HESA, 2022; Advance HE 2023), there is a clear need for subject-specific approaches to EDI support and provisioning.

One approach to challenging lack of EDI support is embedding an ‘inclusive leadership model’. Crucial to any academic environment is strong and responsive leadership, and leaders are key for facilitating and promoting inclusion. An emerging approach to building inclusive research environments is to foster inclusive and departmental-specific leadership skills across all job families, regardless of seniority. By recontextualising typical leadership styles through an inclusive lens, inclusive leadership approaches can challenge slow rates of institutional EDI change and facilitate deeper engagement with changing research cultures (Söderhjelm et al., 2018; Randel et al., 2018).

This article explores the School of Molecular Biosciences’ approach to creating and embedding a holistic approach to inclusive leadership within the department. The outlined Inclusive Leadership Programme aimed to empower all members to consider themselves as leaders, provide a shared definition of inclusive leadership attributes, and embed a baseline of inclusive culture within the School. Here, we present a summary of our approaches and outcomes, as a case study for how other bioscience departments can use co-creation and data-driven EDI approaches to promote whole-school leadership empowerment.

### Identifying Our Inclusive Leadership Traits

The School of Molecular Biosciences (SMB) at the University of Glasgow is a multidisciplinary research and teaching department with a molecular biology focus. Every two years, SMB undertakes a School culture survey. In 2020, respondents highlighted concerns around work/life balance and leadership pressures. Alongside this, our department underwent a significant organisational restructuring process, which provided an opportunity to integrate new EDI initiatives through a whole-School programme. An Institutional Strategic Support Fund was awarded from a centrally held Wellcome Trust Fund to recruit an Inclusive Leadership Coordinator to manage development of the project.

To understand our members’ lived experiences of inclusion and leadership, two co-creation sessions were held online in 2021. The first workshop included PhD students and those with less than eight years’ experience post-PhD, and the second included Technical and Operations staff. We focused these groups’ experiences as they are often excluded from leadership conversations. Participants were asked to explore their own perceptions of what makes an inclusive leader, and reflect on experiences of good leadership, shown in figure one.

### Putting Principles into Practice

Having identified key inclusive leadership traits (Figure 1), a reliable and engaging approach to integrate inclusive leadership into the School was needed. We decided to develop an online learning resource, choosing a platform with interactive content options. Interactive learning significantly improves recall and assists engagement within self-directed learning resources (Chi and Wylie, 2014).

**Figure 1:**
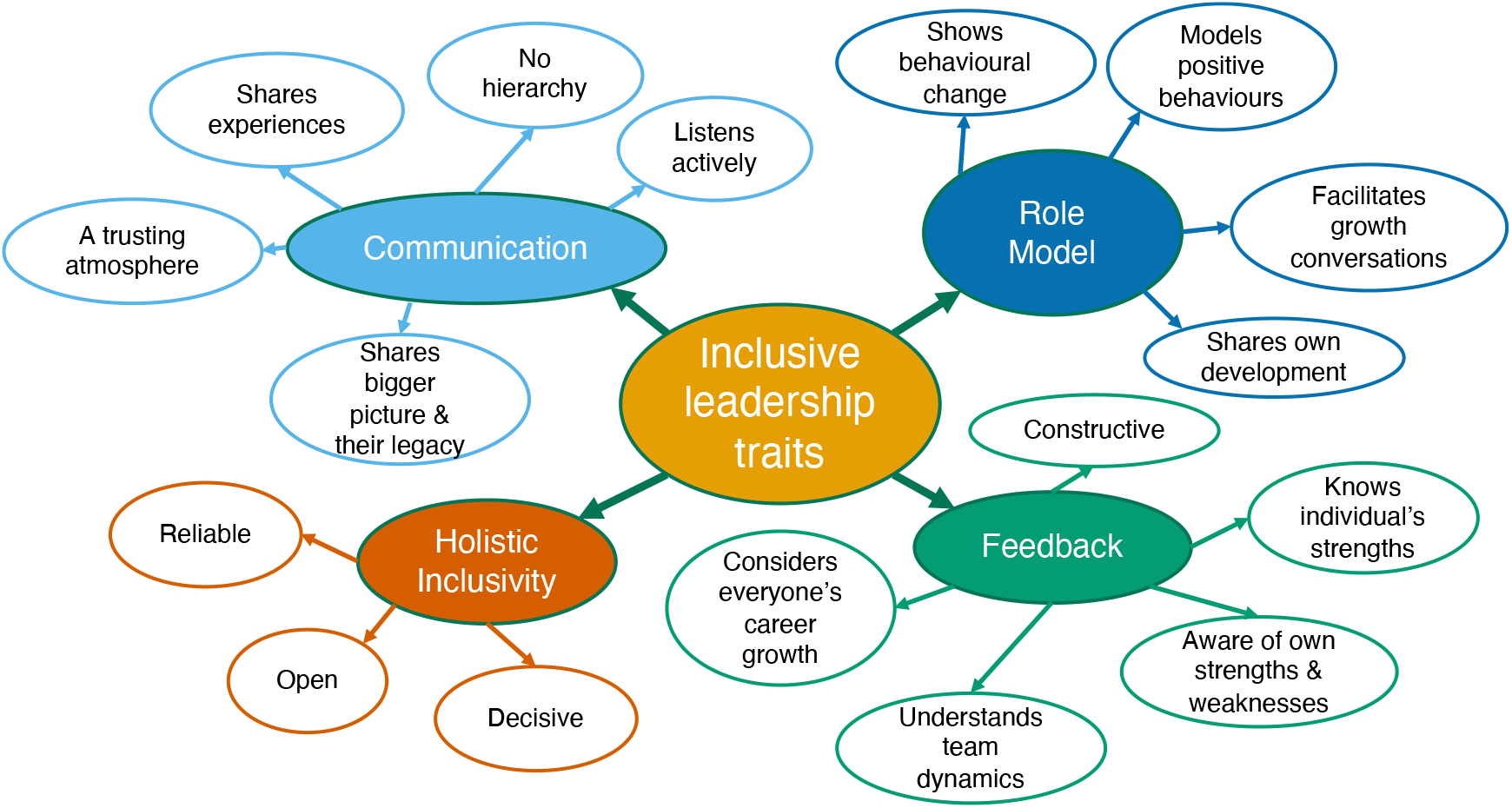
A co-created outline of the School’s identified key inclusive leadership traits. These co-creation identified traits track closely with other examples of best practice inclusivity.

Building on the identified traits an initial draft resource was created. This draft used best practice literature (e.g. Cwik and Singh, 2022; Roberson and Perry, 2022; Booysen, 2013), reported lived experiences, and recorded personal reflections from initial sessions. To refine our self-directed resource, we hosted three co-creation sessions with members of our EDI committee, administrative support team, teaching-focused staff, research-focused staff, PhD students, those <8 years post-PhD completion, and technical and operations staff. These sessions offered space for non-judgemental discussion and peer review, with session content including non-hierarchical language, relevant biosciences examples, lived experiences, and ease-of-use.

On average, the resource takes users between 30 minutes to an hour to complete, with each of the 16 sections taking between two to five minutes. Interactive elements include drop down menus, card sorting, and spaces for self-reflection, illustrated with photos of the school and broader campus. Prompts are provided for participants to further explore their lived experiences and inclusive perceptions. Examples of the resource can be found in Appendix IV.

### Cultivating Support and Integrating Inclusivity

As outlined in the introduction, key drivers for lack of successful EDI integration and uptake are institutionally conservative approaches to balancing cultural adaptation and stability, coupled with a lack of engagement from senior department members. Ensuring a broad uptake and buy-in from all members of the school required communication coordination across the Schools different seminars, group meetings, and internal communication, while building relationships with senior department members.

To build engagement, small information sessions were delivered at senior staff townhall meetings by members of the project, and research students were provided with leaflets at their annual research showcase. Promotion efforts emphasised how ‘everyone is a leader, regardless of their seniority,’ and encouraged members to recontextualise their daily experiences as core parts of inclusive leadership. These sessions were supported with digital posters, reminders in school-wide emails, and with seminars delivered as part of the internal seminar and social series. Those with management responsibilities were encouraged to share the resource with their teaching or research groups.

### Measuring self-reflection and changing behaviours

Unlike many other EDI resources, the Inclusive Leadership resource uses a ‘before’ and ‘after’ scoring system for eight questions (table one). This encourages resource users to consider their self-perceived leadership skills both before and after they have used the resource, providing a gateway to reflective practices while identifying future areas for knowledge building. This also allowed a quantitative understanding of self-perceived inclusive leadership skills, and qualitative investigation of answers provided in free-text sections. Participant demographics are available in Appendix I.

**Table 1:** List of before and after questions used in the Inclusive Leadership Resource.

|  |  |
| --- | --- |
| Communication Section (Figure 2A) | On a scale of 1-5, how do you **currently** rate your **communication skills**? |
|  | On a scale of 1-5, how do you **currently** rate your ability to **present the bigger picture** to your team? |
|  | On a scale of 1-5, how do you **currently** rate your ability to **share your scholarship or research** to your colleagues? |
|  | On a scale of 1-5, how do you **currently** rate your **active listening** skills? |
| Feedback, Recognition, and Development Section (Figure 2B) | On a scale of 1-5, how do you **currently** rate your **ability to manage inclusive meetings**? |
|  | On a scale of 1-5, how do you **currently** rate your overall **feedback, recognition, and development skills**? |
|  | On a scale of 1-5, how do you **currently** rate your ability to **deliver written or oral feedback**? |
|  | On a scale of 1-5, how do you **currently** rate your ability to **encourage personal feedback**? |

Figure 2A shows before and after ratings for communication focused questions. The biggest change in participant’s self-perceived inclusive communication skills is in ‘sharing the bigger picture’, where 46% ranked themselves as ‘4 – good’ or ‘5 – excellent’, increasing to 70% after. The largest positive shift in participant’s self-reflective inclusive communication skills is in active listening. The majority (57%) of participants ranked themselves as ‘3 – okay’ before using the resource. After, the number of participants ranking themselves as ‘4 – good’ and 5 – excellent’ rose to 75%. Participants ranking their active listening skills as ‘excellent’ were all principal investigators or laboratory leaders, with at least 11 years’ experience of acting in a leadership position. Comments included “[I want to develop…] A better understanding of the needs of those with different backgrounds from myself.”

**Figure 2:**
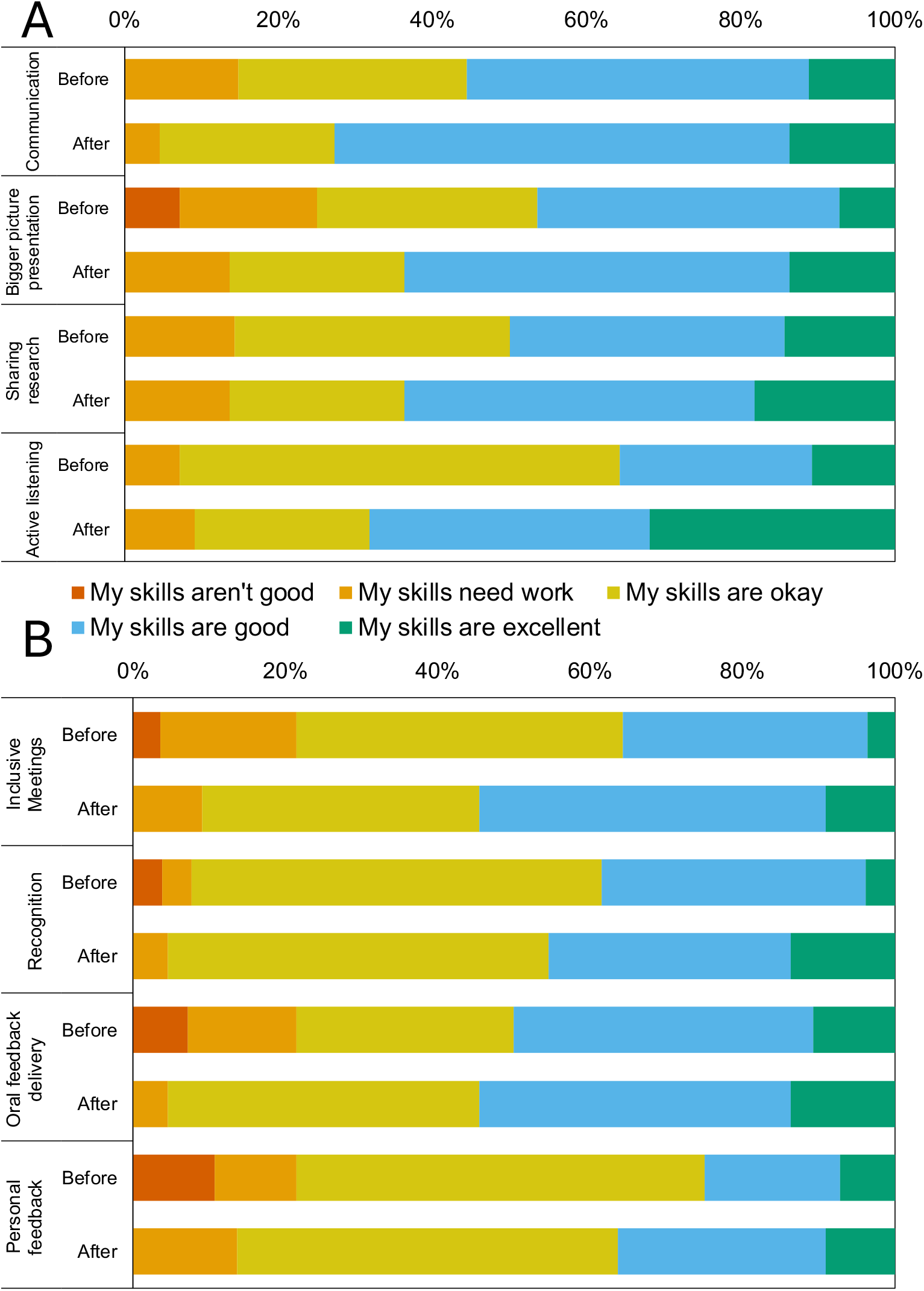
Before and after self-scores considering A) inclusive communication and B) inclusive feedback, recognition, and development. Participants scored themselves from 1-5, where 1 is ‘not good’ and 5 is ‘excellent’. Top row for each question is ‘before’, lower is ‘after’ using the resource.

Figure 2B shows before and after ratings for inclusive feedback, recognition, and development. Of the participants who left text comments about what they’d like to improve, 67% wanted to focus on improving inclusive feedback delivery. The biggest differences in self-scoring before and after using the resource is ‘running inclusive meetings’, where 35% scored themselves ‘4, good’ and ‘5 excellent’ increasing to 60% after using the resource. One participant notes, “[I want to develop…] A better understanding of active listening.” Notably, no participants scored themselves as ‘1 – my skills aren’t good’ after using the resource for any question.

### A reflective evaluation – BOX 1

To understand the impact of the Inclusive Leadership Programme, we undertook discussions with school members involved in co-creation, supporting, and promoting the programme; alongside members who used the resource. These discussions were undertaken by the Inclusive Leadership programme manager, allowing each participant to consider what inclusive leadership means to them, how the programme has impacted their work, and their perceived effect on the school.

Discussions with **The School of Molecular Bioscience’s Head of School**, noted the importance of embedding contextual leadership at an early stage in careers, *“I think leadership should not be seen as something that you just learn on the job, but in my experience a lot of leaders do see it that way*.*”* Helen also highlighted the importance of choice in using resources and training tools, *“One of the things I like about the inclusive leadership as you’ve set it up is that it is not mandatory*… *it is something that is consistently reminded via the newsletter that it is there, and something that the encouragement is to do it repeatedly, to use it as a reflective tool*.*”*

Considering the importance of a bioscience-specific approach, the **School’s Academic EDI lead** adds, *“Whenever I’ve come up seen this kind of reflective approach previously, it’s always bounced off*… *I think having it couched in scientific language and in the lab-based environment is very useful*.*”* He also reflects on the level of information available in the resource, *“If it was any more comprehensive, it would become too overwhelming, I think*… *What we’re trying to do is just increase awareness of another way of doing things*.*”*

These conversations highlighted the importance of engaging with researchers at the beginning of their career, *“This is a good way of developing leadership skills early so PhD students can start thinking about leadership, even when they don’t personally see themselves as a leader yet. If you look at the way you lead on that small scale at the beginning of your career, it allows you to be self-critical and think about how you would do things better in the future. I think in terms of building and developing people into leaders the earlier you start, the better*.*”* **Postdoctoral research associate, EDI team member, and co-creation session participant**.

Equally, the co-creation approach of the resource was highlighted as a key driver for its wide uptake and engagement amongst different members of the School, with a co-creation session participant sharing, *“Firstly, [colleagues] see the effort that’s gone into making the resource, which gets them interacting with it. They can promote it and share, ‘I’ve been involved in this, would you mind completing it*,*’ so you get that extra kind of boost in engagement and participation*.*”* **Postdoctoral research associate, EDI team member, and co-creation session participant**.

### Proactive Inclusivity in Leadership

Reflecting on the Inclusive Leadership Programme as a whole, emerging themes consider the importance of personal proactivity and self-reflection, coupled with a willingness to interrogate pre-formed ideas or beliefs. These traits go hand in hand with inclusivity, and create a foundation for inclusivity as common culture, as opposed to a retroactive addition to a bioscience department’s approaches to EDI.

Of note is the normalisation of lower-end quantitative self-scoring outcomes, where the number of participants ranking themselves ‘1 – not very good’ across the eight questions declined from 11 before using the resource to 0 after. Conversely, those ranking themselves as ‘5 – excellent’ across the eight questions rose from 20 to 27. This indicates a normalisation and reframing of pre-existing skills, providing resource users with a unifying baseline to compare their self-perceived skills to, while considering the success of those skills, and how to improve these skills if necessary.

### Looking Forward: Embedding Inclusivity in Bioscience

The Inclusive Leadership Project will continue to be embedded into the School in new ways. Inclusive Leadership related events run approximately once a term for a variety of different school members, most recently providing space for technicians, operations staff, alongside managerial, professional, and administrative staff. These interactive workshops explore what leadership means to school members and how they can embed inclusivity into their daily work, even if they do not necessarily have leadership aspirations. These discussions provide further opportunity to iteratively update and refine the Inclusive Leadership Resource, accurately reflecting emerging and evolving requirements of the school’s members. This adds a further diversified approach in addition to the original co-creation sessions, improving diversity of experience, equity in access, and inclusion of school members within the process itself.

Creating and assessing EDI-focused whole-School culture change is often challenging, but through continually re-evaluating, promoting, and sharing the resource, we have worked to build a baseline understanding of inclusivity within leadership, and hope to see meaningful change reflected in approaches to peer-peer support, managerial decisions, and self-reflection. The project’s success is further bolstered by the UK’s national research funder, UKRI, now embedding EDI in flagship grant applications (UKRI, 2023). This high-level funder commitment to inclusive research and teaching will continue to enhance EDI awareness, engagement, and commitment within the biosciences.

Equally, the Inclusive Leadership Project highlights the importance of using a ‘human’ approach to bioscience EDI initiatives. As useful as top-level EDI and protected characteristic data-driven approaches to EDI are, this must be coupled with a relevant, peer-created, and collective buy-in approach for success. By embedding bioscience specific examples, such as inclusive lab meetings and laboratory practice, a focus on technicians’ experiences, and historical academic hierarchies, this programme and resource help fill our discipline-specific needs.

Of equal importance was the top-down and bottom-up engagement approach. Senior-level buy-in was supported by the Head of School’s endorsement and promotion of the project at senior staff meetings, which was complemented with workshops and talks at whole-School and meetings and events. This has been further bolstered through our Head of School’s continuous inclusive role modelling throughout her tenure, which she explores further in Walden, 2024. Additionally, by supporting research and teaching leaders to work as inclusive role models, we build a continuous culture of reflexivity and improvement. By engaging all School members, regardless of job role or seniority, we ensured a robust uptake and peer-driven usage to help members embed inclusive leadership into their pre-existing leadership skills and help newer members explore and contextualise their own approaches to leadership.

The Inclusive Leadership Project now extends beyond the School of Molecular Biosciences. To date, it has been shared at AdvanceHE’s EDI Conference in 2024, The Society of Experimental Biology’s Annual Conference from 2022-2024, and was also showcased at the American Association for Phytopathology’s Annual Conference in 2023. The success of our programme at the School level has opened opportunities to support other research and teaching organisations to host skill-share workshops and design their own bespoke resources using our co-creation format. The resource is currently being trialled at multiple Higher Education organisations within the UK, and we are excited to see how organisations adapt it for their own needs. Locally, it has been gratifying to see School members encourage new joining members to use the resource, and to see whole teams work through the resource together.

Ultimately, our vision is to embed proactive inclusivity as a key tenet of any research organisation, with the goal to reduce bullying and harassment, improve feelings of belonging, and create cohesive and enjoyable research and learning environments. It is hard to overstate the importance of feedback, the role of autonomy, and the need for continuous improvement in leadership practices.

An overview of how to set your own inclusive leadership programme in a bioscience department is outlined in box two.

#### BOX TWO

setting up an inclusive leadership programme in your department

**Figure 3:**
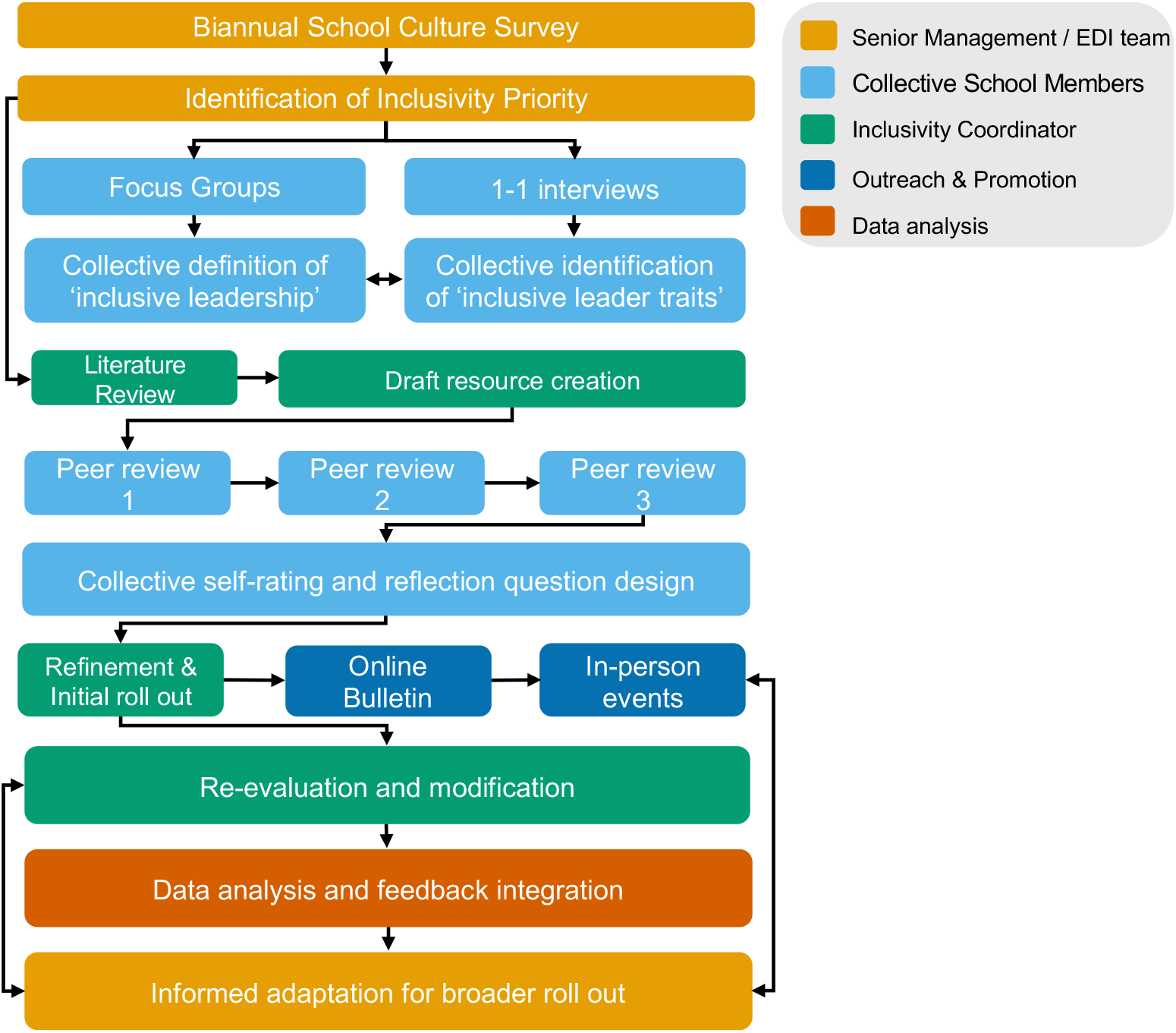
Developing an empowerment and autonomy focused inclusive leadership resource within biosciences departments. Each department comes with their own unique research and experiential landscapes – this framework is not prescriptive, but offers a guide towards iterative creation and resource-buy in. Yellow shows input from senior decision makers, green shows the work of a dedicated inclusivity coordinator, light blue is collective work, dark blue is communication strategies, orange is data analysis requirements.

## APPENDIX I

Information about the resource participants: the number of people they manage, their length of service, and the number of years they have spent in a leadership role (table two). 25 participants provided permission for their anonymous answers to be shared.

**Table S1:** A breakdown of the Inclusive Leadership Interactive Resource participants by the number of people participants have supervisory responsibilities for (top), their current position in the School (middle), and the total number of years they’ve worked.

| Number of people you work with directly or have supervisory responsibilities for | Number of Participants |
| --- | --- |
| None | 2 |
| 1 to 3 | 11 |
| 4 to 6 | 5 |
| 6+ | 7 |
| Current position in the School | Number of Participants |
| Early career researcher (e.g. PhD student, Postdoc) | 8 |
| Principal investigator/lab leader | 10 |
| Technical and Operations | 1 |
| Lecturer, Teaching & Scholarship | 3 |
| Administration & Managerial | 0 |
| Other | 3 |
| Number of years worked in a leadership role. | Number of Participants |
| 1 to 5 | 9 |
| 6 to 10 | 9 |
| 11 to 15 | 4 |
| 16+ | 3 |

## APPENDIX II

List of what participants would like to gain from using the resource

**Table S2:** A snapshot overview of resource participant’s reflections on what they would like to gain from using the resource.

| Respondent number | Respondent comments to: "What would you like to gain from this resource?" | Respondent Number | Respondent comments to: "What would you like to gain from this resource?" |
| --- | --- | --- | --- |
| 1 | always take into consideration other people's perspective | 13 | better awareness of others |
| 2 | Ability to give feedback that's positive and negative with a view to improvement | 14 | Learn to understand my blind spots |
| 3 | communication skills | 15 | Make meetings accessible |
| 4 | Learning to work better with people who think or approach things differently to me, particularly with respect to neurodivergent thought processes | 16 | managing inclusive meetings |
| 5 | Deal with difficult conversations in the lab | 17 | A better understanding of the needs of those with different backgrounds from myself |
| 6 | I would like to be better at presenting my big picture plans | 18 | An approach to help me support the development of people I lead |
| 7 | Different approaches to communication. | 19 | clarity of planning |
| 8 | Giving feedback in inclusive ways | 20 | communication |
| 9 | Feedback skills | 21 | managing group meetings in an inclusive way |
| 10 | Leading meetings and giving feedback. | 22 | As above: the ability to self-promote in an engaging, inclusive manner. |
| 11 | managing expectations | 23 | Listening skills |
| 12 | I know I lack time management (often meetings with team members are longer than they need to be) and assertiveness. | 24 | My ability to recognise people's struggles and points of contact in order to help them best |

**Table S3:**
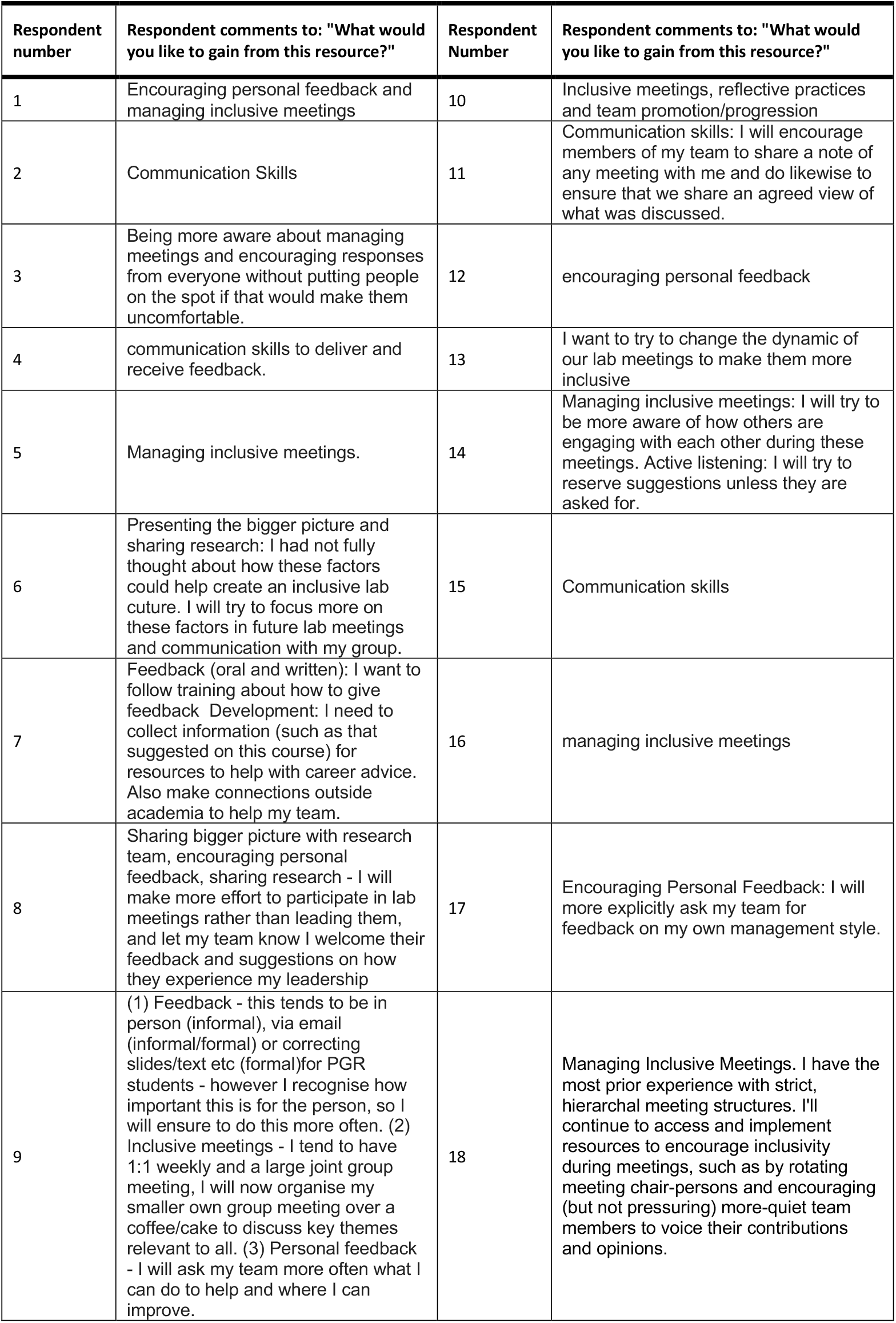
A snapshot overview of what participants would like to work on after using the interactive resource. Note, before and after numbering is not participant-matched.

| Respondent number | Respondent comments to: "What would you like to gain from this resource?" | Respondent Number | Respondent comments to: "What would you like to gain from this resource?" |
| --- | --- | --- | --- |
| 1 | Encouraging personal feedback and managing inclusive meetings | 10 | Inclusive meetings, reflective practices and team promotion/progression |
| 2 | Communication Skills | 11 | Communication skills: I will encourage members of my team to share a note of any meeting with me and do likewise to ensure that we share an agreed view of what was discussed. |
| 3 | Being more aware about managing meetings and encouraging responses from everyone without putting people on the spot if that would make them uncomfortable. | 12 | encouraging personal feedback |
| 4 | communication skills to deliver and receive feedback. | 13 | I want to try to change the dynamic of our lab meetings to make them more inclusive |
| 5 | Managing inclusive meetings. | 14 | Managing inclusive meetings: I will try to be more aware of how others are engaging with each other during these meetings. Active listening: I will try to reserve suggestions unless they are asked for. |
| 6 | Presenting the bigger picture and sharing research: I had not fully thought about how these factors could help create an inclusive lab culture. I will try to focus more on these factors in future lab meetings and communication with my group. | 15 | Communication skills |
| 7 | Feedback (oral and written): I want to follow training about how to give feedback Development: I need to collect information (such as that suggested on this course) for resources to help with career advice. Also make connections outside academia to help my team. | 16 | managing inclusive meetings |
| 8 | Sharing bigger picture with research team, encouraging personal feedback, sharing research - I will make more effort to participate in lab meetings rather than leading them, and let my team know I welcome their feedback and suggestions on how they experience my leadership | 17 | Encouraging Personal Feedback: I will more explicitly ask my team for feedback on my own management style. |
| 9 | (1) Feedback - this tends to be in person (informal), via email (informal/formal) or correcting slides/text etc (formal)for PGR students - however I recognise how important this is for the person, so I will ensure to do this more often. (2) Inclusive meetings - I tend to have 1:1 weekly and a large joint group meeting, I will now organise my smaller own group meeting over a coffee/cake to discuss key themes relevant to all. (3) Personal feedback - I will ask my team more often what I can do to help and where I can improve. | 18 | Managing Inclusive Meetings. I have the most prior experience with strict, hierarchal meeting structures. I'll continue to access and implement resources to encourage inclusivity during meetings, such as by rotating meeting chair-persons and encouraging (but not pressuring) more-quiet team members to voice their contributions and opinions. |

## APPENDIX III

Significant changes by question before and after using the resource

**Figure S4:**
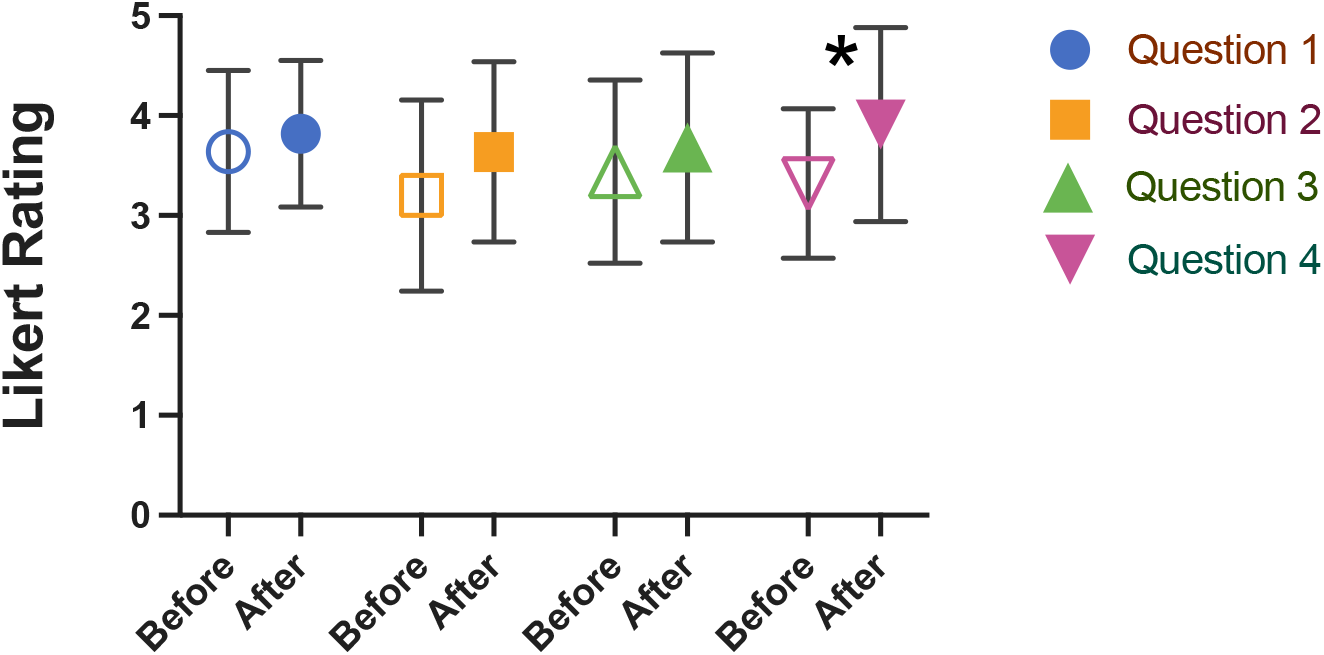
Average self-ranking scores for resource participants before using the resource (open symbols), and after (closed symbols), considering inclusive communication. For question list see Table 1. Averages are transformed Likert scale scores. n=28, p<0.001, Welch’s two sample t-test, bars are standard deviation.

**Figure S5:**
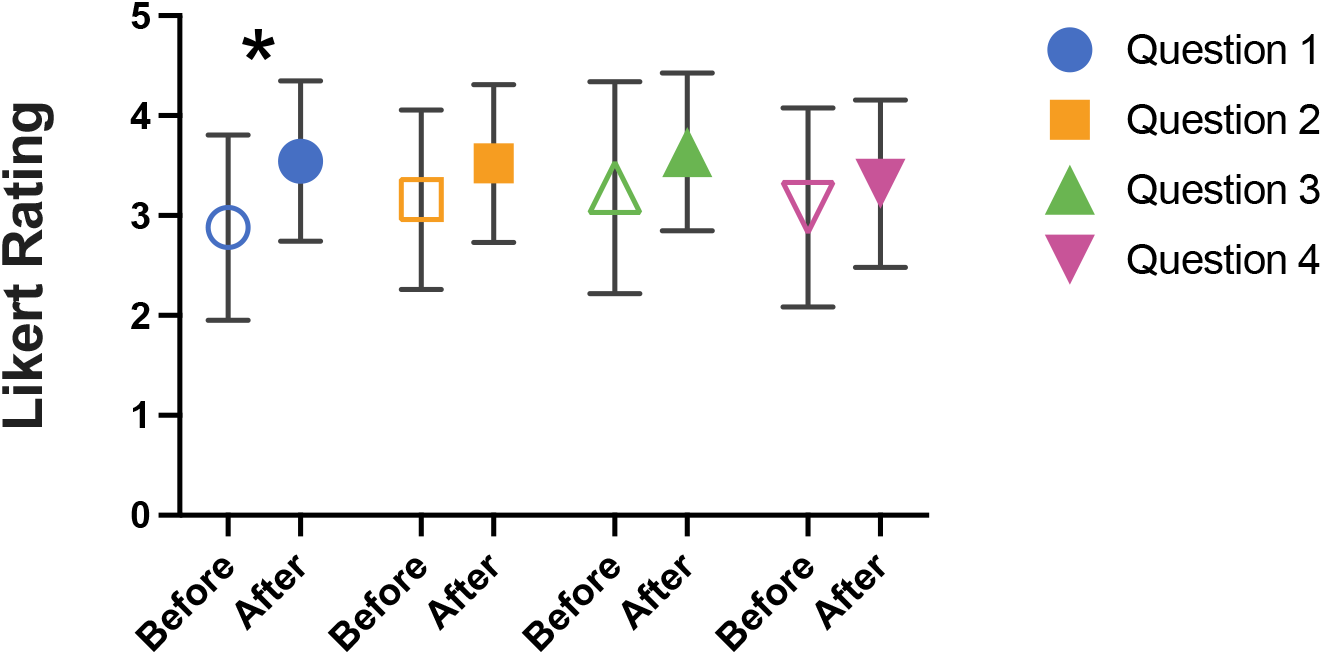
Average self-ranking scores for resource participants before using the resource (open symbols), and after (closed symbols), considering inclusive feedback, recognition, and development. For question list see Table 1. Averages represented transformed Likert scale scores. n=28, p<0.001 using Welch’s two sample t-test, bars are standard deviation.

## APPENDIX IV

Screengrabs of the Inclusive Leadership Resource

**Figure S3:**
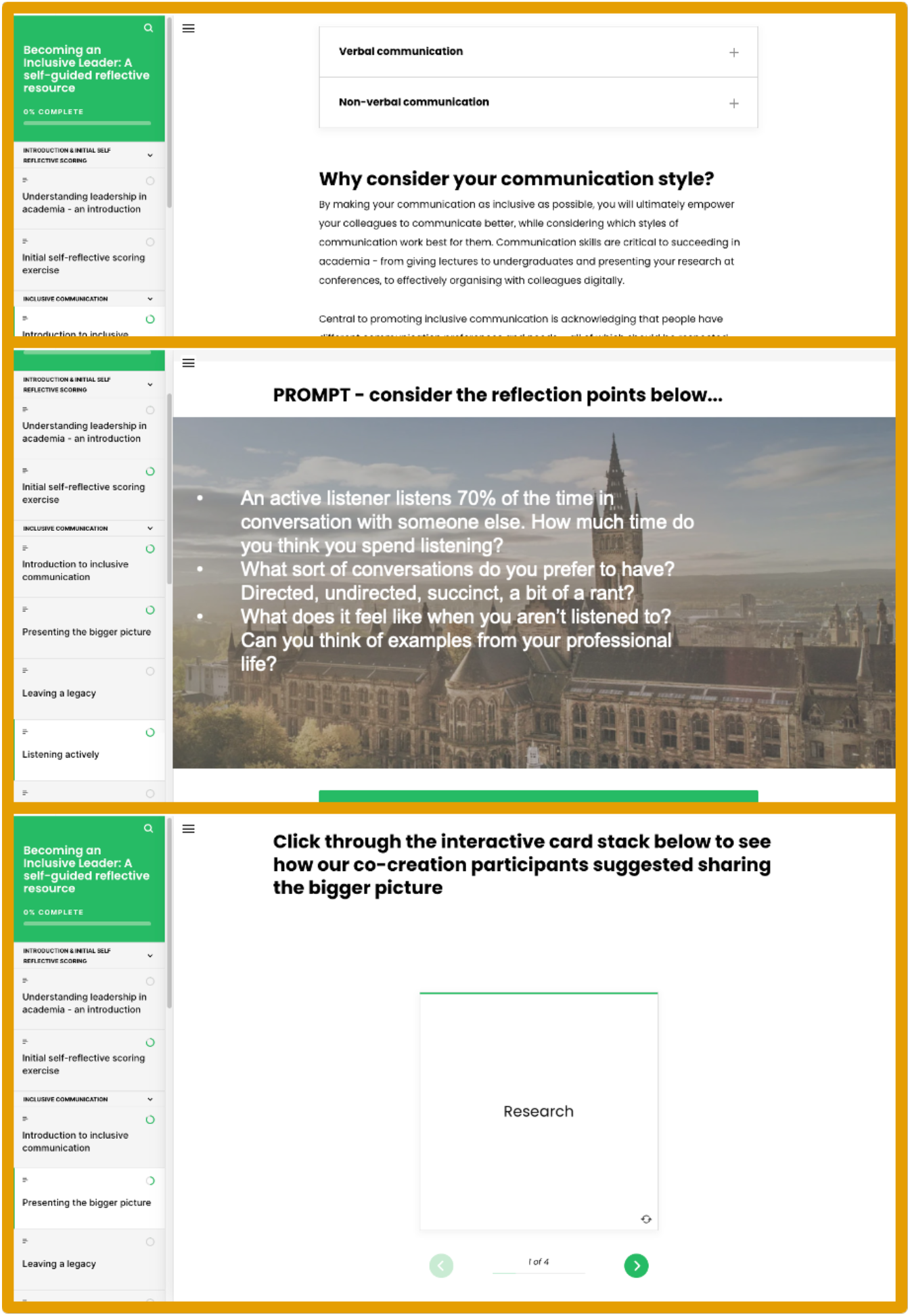
Screengrabs from the Inclusive Leadership Resource considering communication style (top), active listening (middle), and sharing the bigger picture (bottom)

## Notes

### Competing Interest Statement

The authors have declared no competing interest.

